# Oscillatory activity in alpha/beta frequencies coordinates auditory and prefrontal cortices during extinction learning

**DOI:** 10.1101/2020.10.30.362962

**Authors:** Howard J. Gritton, Jian C. Nocon, Nicholas M. James, Eric Lowet, Moona Abdulkerim, Kamal Sen, Xue Han

## Abstract

Cortical synchrony is theorized to contribute to communication between connected networks during executive functioning. To understand the functional role of neural synchrony in cognitive flexibility, we recorded from auditory cortex (AC) and medial prefrontal cortex (mPFC), while mice performed an auditory extinction learning task. We found that while animals gradually showed reduced responding to the unrewarded tone over hundreds of trials, the power of local field potential (LFP) oscillations (8-18 Hz, centered at alpha/beta frequencies) in AC and mPFC exhibited immediate and robust increases, prior to behavioral changes. The strength of LFP alpha/beta power in the mPFC, but not AC, was strongly correlated with the behavioral performance that mice would achieve later in the training session. Further, we found that coherence between AC and mPFC at 8-18Hz was selectively enhanced only after mice learned to suppress licking, and this LFP coherence increase coincided with a reduction in spiking rate for the unrewarded tone in AC. These results reveal that enhanced interactions between PFC and AC is an inherent property of auditory discrimination learning, and that coordinated alpha/beta oscillations contribute to cognitive flexibility.

## Introduction

The ability to adapt to ever-changing environments represents a key tenet of cognitive function and executive control. Fundamental to this process is the integration of sensory cues, decision making, and the executive control of behavior (Miller and Cohen, 2001; Dalley et al., 2004). Deficits in executive function also represent a hallmark of many psychiatric illnesses including schizophrenia, ADHD and bipolar disorder (Hiser and Koenigs, 2018). Many higher order cognitive processes, including sustained attention, cognitive flexibility, and behavioral inhibition, are integrally tied to prefrontal networks specifically (Baddeley, 1998; Stuss and Alexander, 2000; Logue and Gould, 2014). The prefrontal cortex (PFC) can be further separated both functionally and anatomically into the orbital prefrontal cortices, and the medial prefrontal cortex (mPFC) which is closely tied to attentional processing and behavioral flexibility. The ventral mPFC consists of the more anterior prelimbic cortex and the more ventral infralimbic cortex. The dorsal mPFC includes the precentral cortex and anterior cingulate cortex. Lesions of the anterior cingulate impede associational learning and working memory tasks, particularly if they are associated with motor control (Rushworth et al., 2003). mPFC neurons show a high degree of selectivity for conditioned sensory stimuli and can be differentially regulated by context or task demands (Maren and Quirk, 2004; Chang et al., 2010; Euston et al., 2012; Hyman et al., 2012; Moorman and Aston-Jones, 2015). Along the dorsal/ventral axis of the mPFC, a dichotomy also exists between function. Prelimbic cortex regions respond to predictive sensory cues and are thought to promote cue mediated behaviors including reward seeking. In contrast, infralimbic regions are thought to be essential for behavioral inhibition, in particular stop related signals associated with extinction learning (Barker et al., 2014; Gourley and Taylor, 2016).

Extinction learning occurs when a well learned association between paired stimuli ceases to be conditionally reinforced, and the subject shows a diminished behavioral response. The mPFC not only responds to conditioned cues but plays a causal role in the extinction of associative memories in rodents (Van den Oever et al., 2013). The cortical circuit mechanisms for behavioral inhibition during extinction learning are not completely understood, but it has been hypothesized to involve diminished sensitivity for previously conditioned cues in sensory processing regions. It is also possible that changes in responsivity for sensory cues could involve periods of enhanced connectivity between prefrontal and sensory regions during stimulus processing, as cross-regional coherence is theorized to contribute to communication between brain regions during cognition.

To better understand how sensory and prefrontal regions interact during extinction learning, and if functional coherence between these brain structures contributes to cognitive flexibility, we performed simultaneous recordings from auditory cortex (AC) and mPFC while mice underwent an auditory extinction learning task. We found that an increase in alpha/beta frequency (8-18Hz) LFP power in the mPFC and AC as extinction training began, and that changes in LFP alpha/beta frequency oscillations in mPFC, but not AC, were predictive of future performance. We also observed that coherence between PFC and AC at these frequencies strengthened as extinction learning progressed, and the occurrence of maximal coherence coincided with a reduction in AC neural firing rates to the now unrewarded tone. Together, these results suggest that changes in alpha/beta frequency power emerges in mPFC and AC as task rules are updated, and the change in mPFC is selectively involved in the suppression of previously learned behaviors in a way that informs future performance. Furthermore, the coordination of activity in LFP alpha/beta frequencies between PFC and AC could contribute to the strength of responding seen in sensory region important for processing sensory stimuli.

## Results

### An auditory extinction task designed to examine behavioral flexibility

Although it is well known that mPFC is involved in cognitive flexibility, it is less well understood how the representations of stimuli in primary sensory regions are modified when task rules are updated, and if executive regions contribute to these changes. In order to understand how mPFC and AC interact during a change in task rules, we simultaneously recorded from both structures using multi-contact electrodes as animals underwent extinction training for a previously rewarded tone (Figure 1C). Mice were first trained over 1-2 weeks to acquire an operant association between a predictive auditory stimulus and reward availability. Training occurred over several stages with initial habituation to head fixed condition, waterspout acclimation, and learning to lick for a water reward (Figure 1A). Animals were trained on 2 tones separated by ~1.5 octaves (low tone: 3075Hz, or high tone: 10035Hz) – both of which were rewarded (Figure 1B; gray - standard task). A trial was rewarded following a conditioned response (lick to a dry reward port) any time within 2 seconds of cue onset. A reward consisted of delivery of 5μL saccharin sweetened water. Prior to the extinction session, animals were performing more than 350 rewarded trials with a response rate greater than 80 percent during a 60 min training session. On the extinction day, animals underwent extinction training for 1 of the 2 tones that had been previously conditioned, while the other tone maintained its reward relationship (Figure 1B, blue). Acute multichannel recordings were performed on the extinction day, where, animals gradually learned through trial and error that licking for the extinguished tone is no longer rewarded.

**Figure 1:**
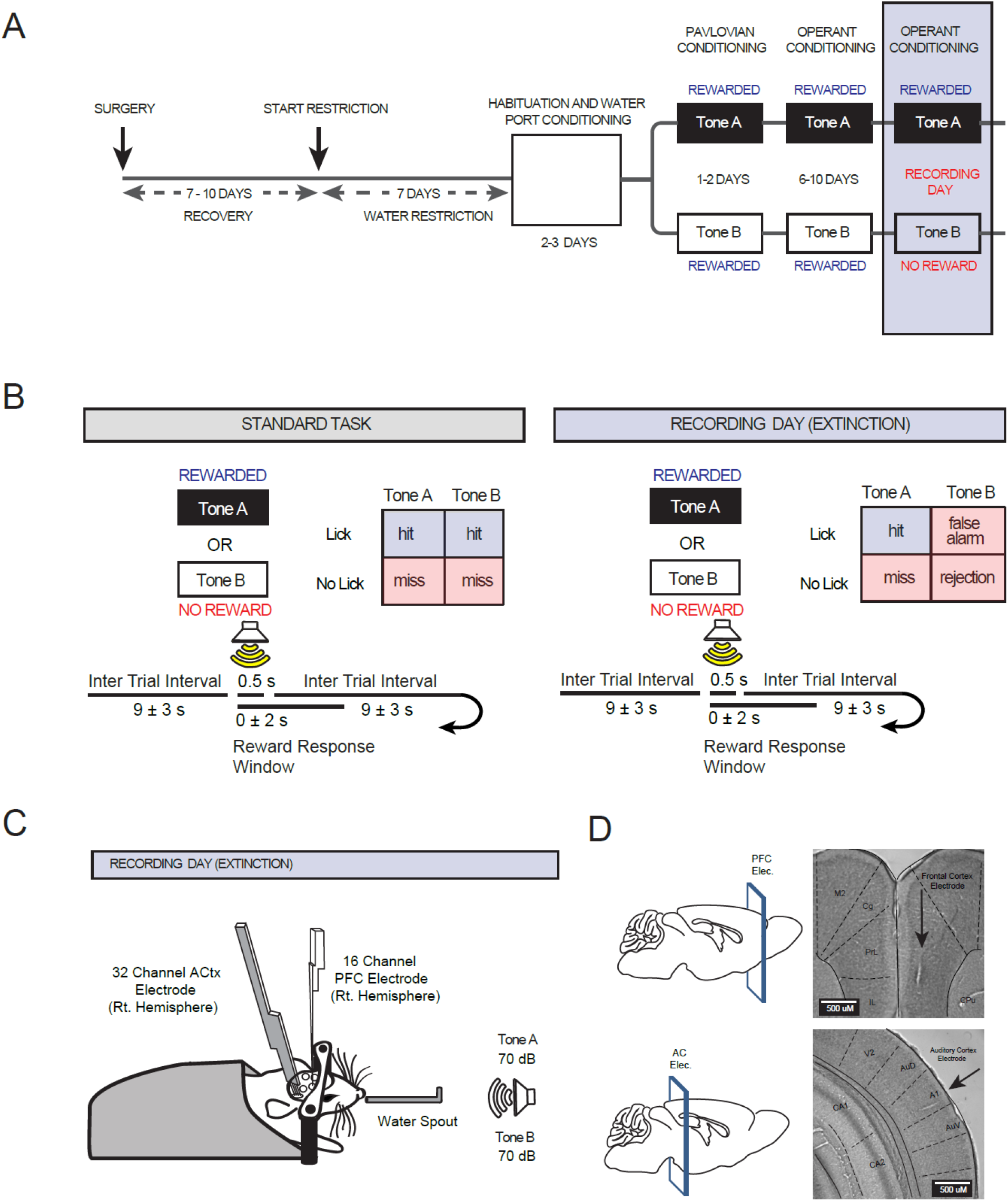
Experimental timeline and behavioral training paradigm. **A:** Mice underwent surgery for head bar fixation and markers for electrode placement. Following recovery and gradual water restriction, animals were trained for 10-14 days in a shaping paradigm to ultimately perform a two-tone operant conditioning task. Animals were trained to criterion performance before the recording day (>80% response to both tones over 350+ trials). On the recording day, electrodes were lowered into PFC and AC. Animals were given 50 standard task trials (both tones rewarded) prior to undergoing extinction training for one of the two tones. **B:** Task conditions and outcomes differed by reward contingencies. (Standard task - Left): Animals were trained to lick for a small water reward following the presentation of a 500ms pure tone. Mice received ~5μL of water if a lick response was recorded for either tone (3.075kHz or 10.035kHz) within 2s of tone presentation on the standard task. Tones not followed by a lick within 2s of tone onset, were unrewarded and were counted as a miss. Tones were presented at random with an inter trial interval of 9±3 s. (Extinction Task - Right): On the recording day, following 50 standard baseline task trials (BASE), animals started the extinction portion of the task where the reward was removed for one of the two tones. The extinguished tone (3.075kHz or 10.035kHz) was randomized across subjects. During extinction, licks for the non-rewarded tone were counted as false alarms (FA’s) while suppression of the licking response was counted as a correct rejection (CR). Hits and misses were counted for the still rewarded tone as previously described. **C:** Head fixed mouse preparation with electrode placements during the recording day. One 16-channel electrode was positioned in medial prefrontal cortex (PFC) and a 32 channel, 4 shank, recording electrode was targeted to auditory cortex (AC). PFC single shank penetration occurred along the dorsal-ventral axis with sites facing medially. For the AC array, insertion was at a 30° angle with all recording sites facing medially while the shanks were situated along the rostral-caudal axis. **D:** Cartoon depiction of electrode placement locations with nissl stained histology sections shown to the right indicating electrode placement. Data was collected from 16 animals. **Abbreviations:** Cg; Cingulate Cortex, Prl; Prelimbic cortex, IL; Infralimbic Cortex, CPu; Caudate Putamen, M2; Secondary Motor Cortex, AuD; Secondary Auditory Cortex Dorsal, A1; Primary Auditory Cortex, AC; Auditory Cortex, PFC; Prefrontal Cortex.

### Mice reduce responding for the extinguished tone over several hundred trials in a single extinction training session

We evaluated learning by measuring the relative rate of responding to the still rewarded tone (hit rate), and the strength of behavioral inhibition by responses to the unrewarded tone (false alarm rate). We calculated the sensitivity index or d-prime, as a combined measure of the strength of responding to the continuously rewarded tone relative to the strength of suppression for the no longer rewarded tone. D-prime was calculated as the difference between the hit rate for the rewarded tone and the false alarm rate. For each animal, the d-prime score was calculated on each trial using a sliding window of 50 trials in single trial increments. Figure 2A illustrates the behavioral acquisition curve for a representative animal throughout the recording session.

**Figure 2:**
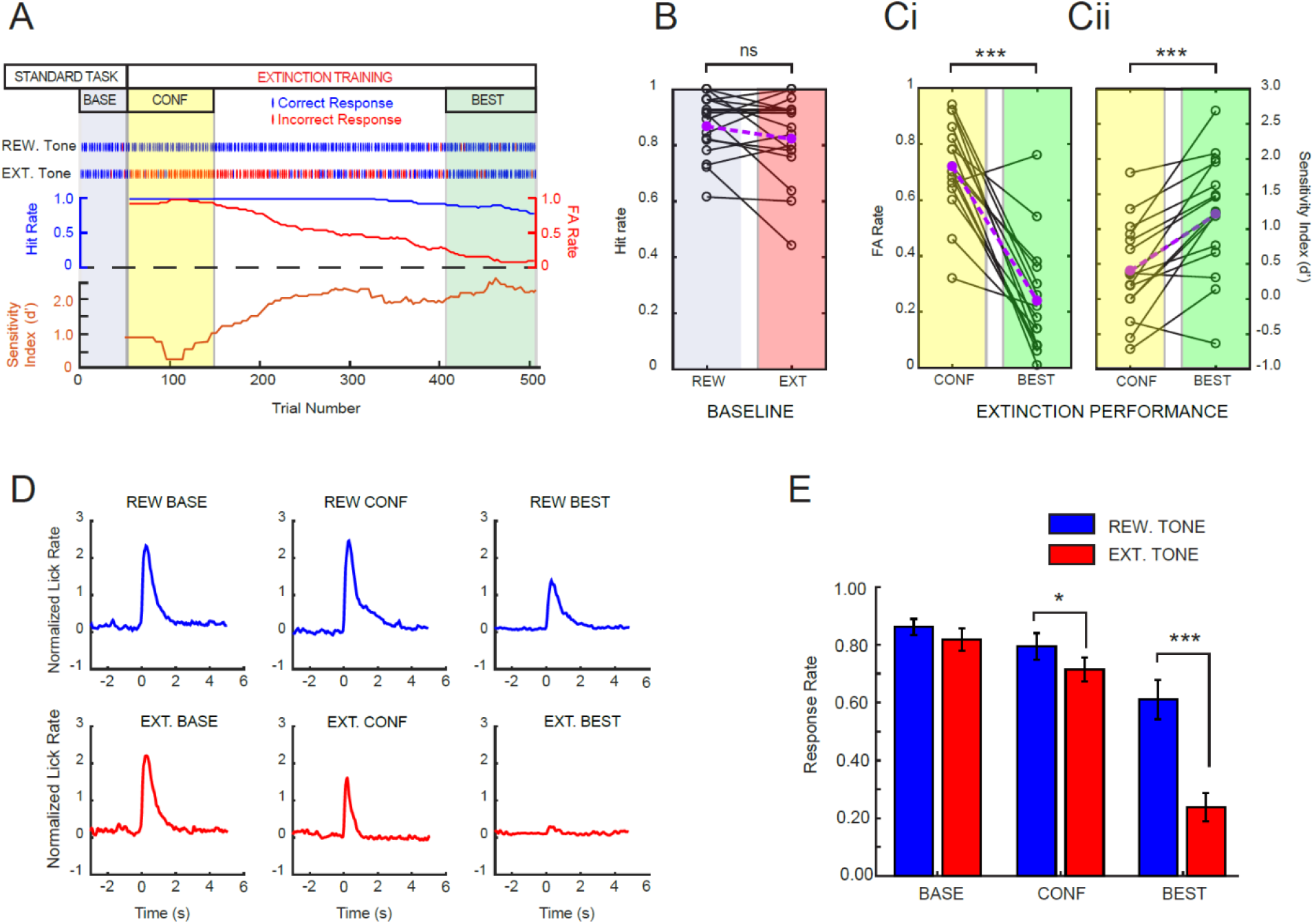
Behavioral performance during extinction learning. **A:** Individual session performance from a single mouse from the extinction training session. Performance was compared across 3 stages: a baseline stage (BASE) consisting of 50 standard trials, a confusion stage (CONF: yellow) which represents the first 100 trials after extinction begins, and the 100 best performance trials (BEST: green). For the example mouse, individual trial responses (blue and red ticks) and boxcar avg. hit rate, false alarm rate and a composite score (sensitivity index or d-prime: *d’* = *Z*(hit rate) - *Z*(false alarm rate)) are shown below. Note the reduction in FA rate and the increase in d-prime over training. **B:** Hit rate performance for all 16 animals from the 50 baseline trials across the two tones. Animals show consistent responding to both tones: response rate (> 80%) for both tones and not significantly different from one another (t(15)=1.437; p=0.171). Population average is shown in purple. **C:** False alarm (FA) rate **(i)** and Sensitivity Index (d’) performance **(ii)** for all 16 animals comparing performance between the CONF and the BEST performance stages. Animals showed a significant reduction in FAs (i) from the CONF to the BEST stages (t(15)=7.451; p=2.20e-6). Population average is shown in purple. Sensitivity index (ii) performance showed a significant increase in tone discrimination performance across the two tones between the CONF and BEST stages (t(15)=5.167; p=1.15e-4). Population average is shown in purple. **D:** Normalized lick rate for both tones across three stages of training from all mice. Under the BASE condition both tones are rewarded. During the CONF and BEST stages, EXT tones are no longer rewarded (red). **E:** Quantification of the population response rate (lick trials) in the 0-2 seconds following tone onset showing significant changes in response rate by block (ANOVA(2,15), F=39.755, p=3.673e-9), by tone (ANOVA(1,15), F=23.854, p = 1.98e-4), and a significant tone by block interaction (ANOVA(2,30), F=20.751, p=2.198e-6). Post hoc analysis revealed significant differences between tones in both the CONF (p=0.0466) and the BEST (p=4.231e-5) performance stages. Error bars represent S.E.M. *=pd<0.05, ***=p<0.001.

On the extinction training day, mice first received 50 standard conditioning trials (25 of each tone, the baseline stage) where both tones were rewarded, as in all training sessions on previous days. Hit rate did not differ between the two tones (Figure 2B). The extinction stage began on trial 51 where the reward contingency was removed from one of the two tones and continued until responding for the still rewarded tone fell to below 60% in the 50-trial moving average. The extinguished tone (EXT Tone, high or low frequency) was counterbalanced across animals. Because mice learn at different rates, we drew comparisons between three intervals, the first 50 trials during the baseline (BASE) stage, the first 100 extinction trials following the removal of reward that we termed the confusion (CONF) interval, and the 100 consecutive extinction trials during the remainder of the training session where the animal reached their best performance (BEST) as defined by highest d-prime scores. Animal reached their best performance on average at 188.125 ± 18.58 rewarded trials or a total of 566.76 ± 74.8 trials (mean ±S.E.M, n=16 mice: note that no more than half of the extinction trials can be rewarded) We found that 14 of 16 subjects showed marked improvement across the training session, with significant reductions in false alarm rates (Figure 2Ci) and significant increases in their d-prime scores (Figure 2Cii). In addition, mice generally showed a higher lick rate for rewarded tones at the beginning of training sessions when thirst is greatest relative to the middle or end of the session, even though correct response rates remained high and misses were low. When we compared the response rate during extinction, we found a small but significant difference in response rates (8.00±3.69, mean ±S.E.M) between the two tones for the confusion stage, and the difference became much greater (37.44±6.57, mean ±S.E.M) for the best performance stage (n=16 mice, figure 2E).

### mPFC LFP power is selectively modulated at gamma frequencies versus alpha/beta frequencies during different stages of extinction training

In order to investigate how extinction training influences cortical changes during behavioral performance, we compared the changes in mPFC LFP power to the two tones during the baseline stage (both rewarded, Figure 3A and Figure 3Bi), during the confusion stage, and the best performance stage (Figure 3 Bii-Biii). During the baseline stage, when both tones are still rewarded, it is evident that the tone produced robust changes in LFPs. In general, the sound evoked related potentials (ERPs) were evident ~30 ms after tone onset. Tone presentation produced a transient broadband increase in power at frequencies above ~30Hz, and a strong and prolonged reduction in low frequency power (0.5-20Hz). This reduction in low frequency power coincided with the period of reward retrieval that contained active motor licking responses beginning at ~100ms after tone onset.

**Figure 3:**
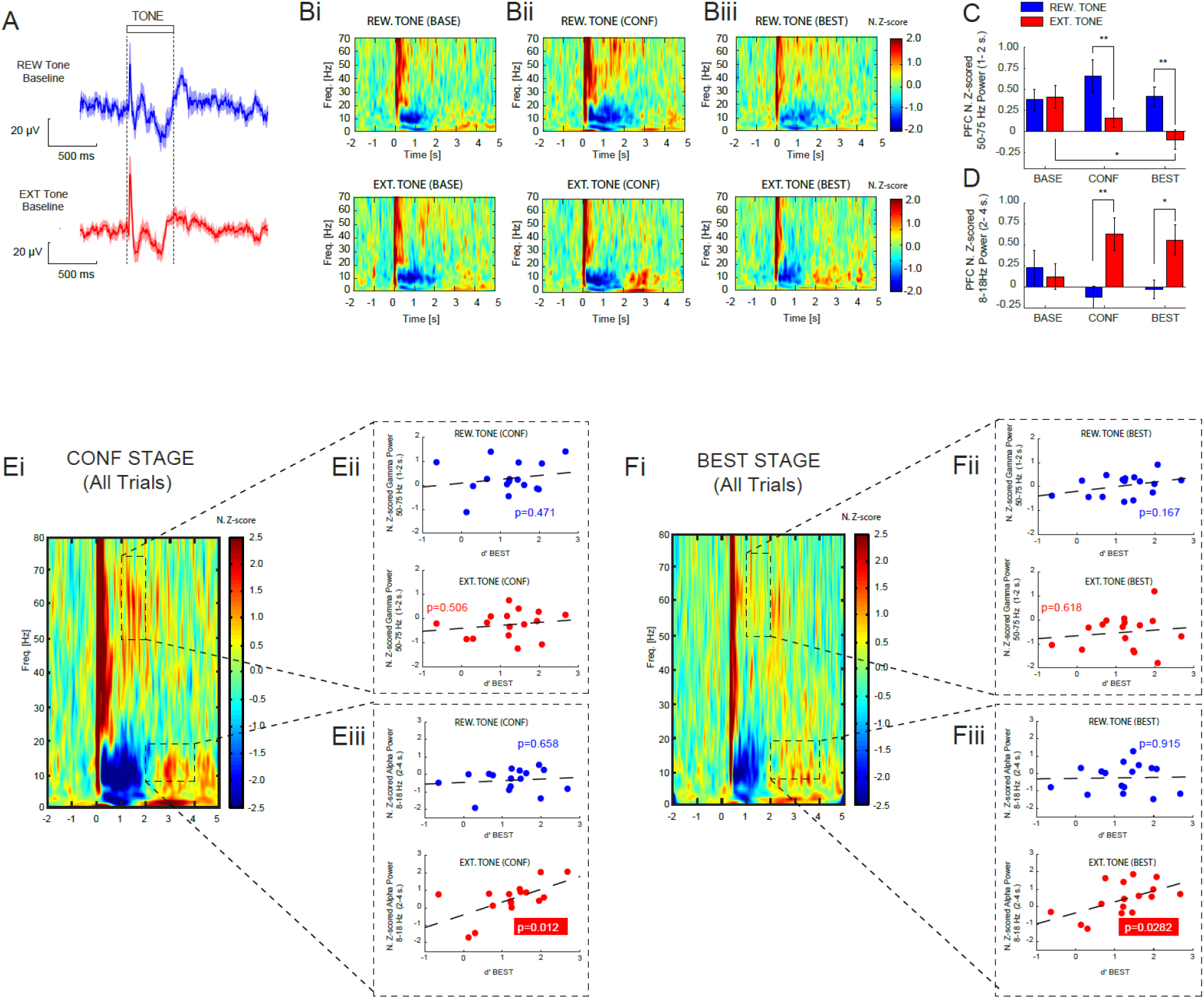
Alpha/beta and gamma power in the PFC discriminate trial types during extinction training and correlate with the learning rate. **A:** Mean population baseline ERP local field potential (LFP) response from all PFC channels during tone presentation for the two tones (REW:top and EXT:bottom) prior to extinction training for all animals. **B:** Average normalized wavelet spectrogram of PFC LFPs aligned to tone onset during the BASE **(i)**, CONF **(ii)**, and BEST **(iii)** performance stages from all mice for the rewarded tone (top) and the extinguished tone (bottom). Tone onset begins at zero. Power was Z-score normalized to the 2s window before tone onset. **C:** Quantification of changes in gamma power (50-75 Hz) in the 1-2 second reward window following tone onset across the three stages of training. Gamma showed significant changes by tone (ANOVA(1,15), F=12.457, p=0.003) and a significant tone by block interaction (ANOVA(2,30), F=4.444, p=0.020). Post hoc analysis revealed significant differences between tones in both the CONF (p=0.004) and the BEST (p =0.01) performance stages while there was a significant reduction in gamma power for the EXT tone between the BASE and BEST performance stages (p=0.0315). **D:** Quantification of changes in alpha/beta power (8-18 Hz) in the 2-4 second post-reward window across the three stages of training. Alpha showed significant changes by tone (ANOVA(1,15), F=9.584, p=0.007) and a significant tone by block interaction (ANOVA(2,30), F=5.657, p=0.008). Post hoc analysis revealed significant differences between tones in both the CONF (p=0.003) and the BEST (p=0.0243) performance stages. **E:** Average normalized wavelet spectrogram from the PFC for all CONF trials **(i)** and relationship of gamma power **(ii)** and alpha/beta power **(iii)** on future performance for each tone (blue:REW tone and red:EXT tone). Regression analysis compares d’ from BEST performance period to gamma (ii) or alpha/beta (iii) power from confusion interval. There was a significant correlation between the strength of alpha/beta power following EXT tone presentation on CONF trials when correlated to asymptotic d’ performance (R=0.611, p=0.012, n = 16 mice). **F:** Average normalized wavelet spectrogram from the PFC for all BEST trials (i) and relationship of gamma power **(ii)** and alpha/beta power **(iii)** on performance for each tone (blue:REW tone and red:EXT tone). Regression analysis compares d’ from BEST performance period to gamma (ii) or alpha/beta (iii) power from the BEST interval. There was a significant correlation between the strength of alpha/beta power following EXT tone presentation and d’ score during the BEST performance block (R=0.547, p=0.028, n = 16 mice). Error bars represent S.E.M. (*=p<0.05, **=p<0.01).

To determine what LFP changes were related to motor activity that are not primarily related to the cognitive aspects of the task, we analyzed spontaneous licking behavior during inter-trial intervals (ITIs). Because mice periodically licked the reward port throughout the training sessions, including ITIs, we randomly selected 50 spontaneous licks that occurred during the ITI from each animal, and compared the LFP spectral changes under those conditions to the tone-evoked LFP changes. During spontaneous licking, there was a robust and prolonged decrease in low frequency power (figure S1). This reduction in low frequency power (0.5-20Hz) during licking following the tone was consistently present across all blocks of trials including baseline, confusion, and best performance stages, similar to that detected during spontaneous licking during the ITI. This suggests that the prolonged reduction in LFP low frequency power after tone onset is mainly a result of motor execution, unrelated to the cognitive aspects of the task (Figure S1). This is consistent with reports of decreases in beta band power during motor execution in rodents, monkeys, and humans (Nikouline et al., 2000; Pfurtscheller et al., 2003; Tzagarakis et al., 2010; Park et al., 2013).

When we focused on LFP changes unique to the confusion stage, we found that there were significant reductions in the higher gamma frequency (50-75 Hz) power for the EXT tone relative to the still rewarded tone, during the reward retrieval window (1.0-2.0s, Figure 3Bii). The gamma power difference between the two tones arises from reduced gamma power for the EXT tone and elevated power for the REW tone (Figure 3C). Gamma oscillations have been shown to be synchronized between cortical areas during selective attention (Buschman and Miller, 2007; Gregoriou et al., 2009; Hipp et al., 2011), and auditory evoked gamma power in AC can predict associative learning and the strength of auditory remapping during task acquisition (Headley and Weinberger, 2011). Thus, the observed gamma oscillations selectively associated with still rewarded stimuli, may be involved in the execution of cue guided behavior coincident with discrimination performance. In addition, we noticed during the confusion stage, 8-18Hz LFP power was increased during the post-reward response window (2.0-4.0s) specifically for the EXT tone (Figure 3Bii), and primarily driven by augmented power following the EXT tone (Figure 3D). During this confusion stage, changes in LFP power should most strongly reflect changes about the stimulus contingency, as responding for the EXT tone is still largely maintained (~77% false alarm rate). The LFP changes observed in the confusion stage: increases in gamma frequency power during the reward window for the REW tone, and the reduction in alpha/beta frequency power during the post-reward window for the EXT tone, were maintained into the best performance stage (Figure 4Biii). Together, these results demonstrate that increases in mPFC LFP oscillatory power across different frequencies during extinction training arise at different time points following tone presentation and likely serve distinct functional processes related to the regulation of behavioral actions or changes in expectation.

**Figure 4:**
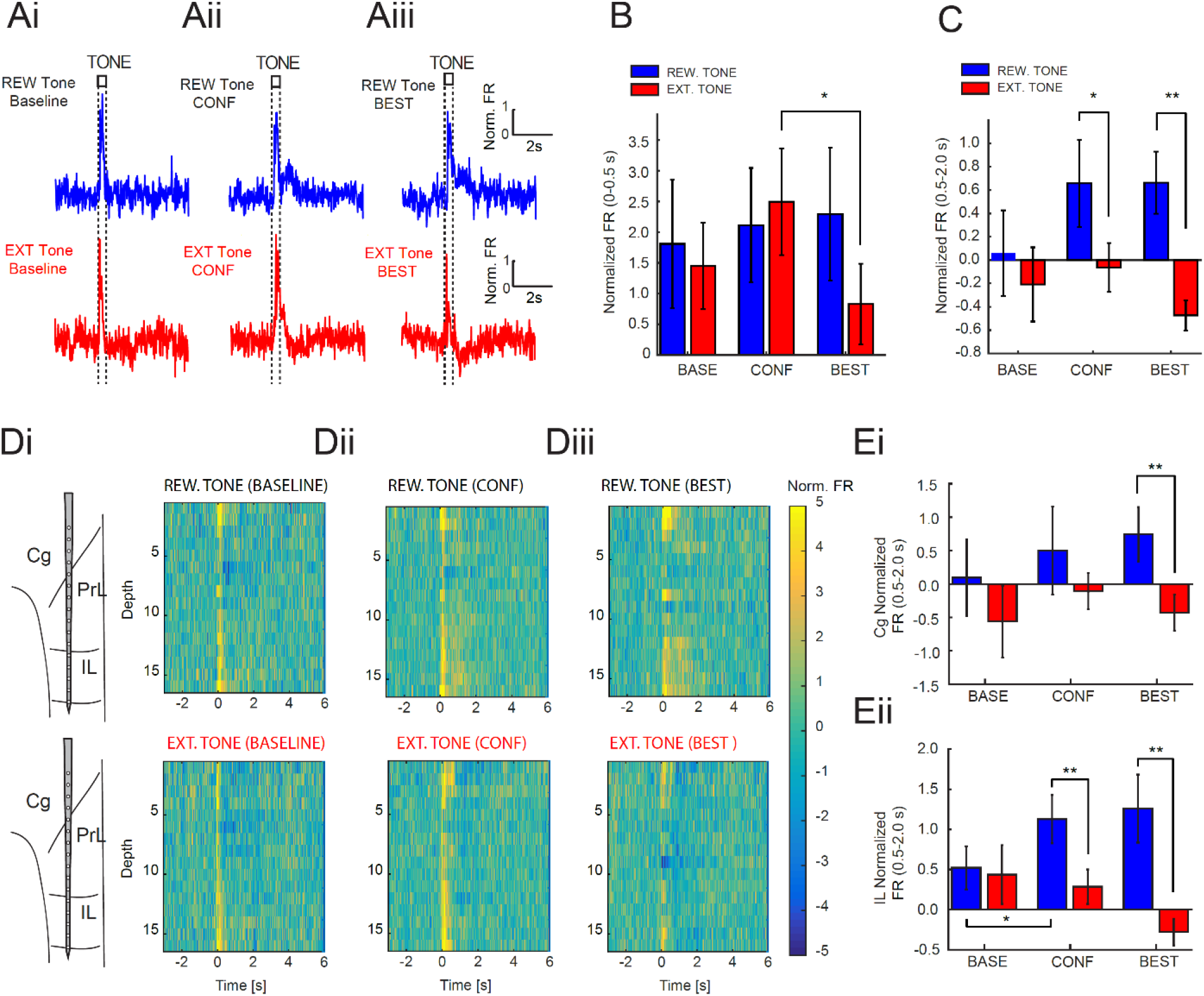
Changes in neuronal firing rate across regions of the PFC encode changes in reward contingencies. **A:** Average sound evoked PFC multi-unit activity (MUA) responses aligned to tone onset during the BASE **(i)**, CONF **(ii)**, and BEST **(iii)** performance stages from all mice for all PFC channels separated by tone: rewarded tone (top:blue) and the extinguished tone (bottom:red). **B:** Quantification of changes in normalized firing rate (FR) during the tone interval (0-0.5s) across the three performance stages. The FR was significantly modulated by an interaction between block and tone (ANOVA(2,30), F=4.0563, p=0.027) and post-hoc analysis revealed a significant reduction in amplitude for the extinguished tone between the CONF and the BEST performance stages (p=0.031). **C:** Quantification of changes in normalized firing rate (FR) during the lick response window (0.5-2.0s) across the three performance stages. MUA was significantly modulated by tone (ANOVA(1,15), F=13.864, p=0.002). Post hoc analysis revealed significant differences between tones in both the CONF (p=0.039) and the BEST (p=0.003) performance stages. Note that the FR is enhanced during the response window following the rewarded tone and reduced for the extinguished tone. **D:** Changes in normalized firing rate in PFC by subregion across the BASE **(i)**, CONF **(ii)**, and BEST **(iii)** performance blocks. Channels are oriented by depth with superficial channels representing cingulate (Cg), intermediate channels representing prelimbic (PrL) regions, and deeper channels representing infralimbic (IL) regions. **E.** Quantification of changes in normalized firing rate (FR) during the lick response window (0.5-2.0s) in cingulate (CG) cortex **(i)** and infralimbic (IL) cortex **(ii)**. MUA was significantly modulated by tone (ANOVA(1,15), F=10.598, p=0.005) in cingulate cortex **(i)**. Post hoc analysis revealed significant differences between tones in the BEST (p = 0.003) performance stage. Infralimbic cortex showed a strong reward outcome sensitivity with MUAs significantly modulated by tone (ANOVA(1,15), F=15.35, p=0.001) and a significant tone by block interaction (ANOVA(2,30), F=4.385, p=0.002: **ii**). Post hoc analysis further revealed significant differences between tones in both the CONF (p=0.004) and BEST (p=0.008) performance stages and enhanced responsivity to the REW tone between BASE and CONF (p=0.038) performance stages. (Cg; Cingulate Cortex, Prl; Prelimbic cortex, IL; Infralimbic Cortex, MO; Medial Orbital Cortex). Error bars represent S.E.M. (*=p<0.05, **=p<0.01). **Abbreviations:** Cg; Cingulate Cortex, Prl; Prelimbic cortex, IL; Infralimbic Cortex.

### mPFC LFP 8-18Hz (alpha/beta) power is correlated with future performance during extinction training

We next examined whether changes in gamma and alpha/beta power from both the confusion and best performance stages are correlated to behavioral performance (Figure 3Ei-Eiii and 3Fi-Fiii). Specifically, we used regression analysis to compare the d-prime behavioral scores from the best performance stage to the relative level of gamma or alpha/beta power either from the confusion stage (Figure 3E) or the best performance stage (Figure 3F). We found that the strength of alpha/beta power following the EXT tone during the confusion stage (trials 51-150) was a strong predictor of the d-prime performance score during the best performance stage, which typically occurs hundreds of trials later (Figure 3Eiii). This relationship was not apparent for the REW tone, or for gamma power for either tone during the confusion stage (Figure 3Eii).

In addition, we further investigated the relationship between gamma (Figure 3Fii) and alpha/beta (Figure 3Fiii) power from the best performance stage to their d-prime behavioral performance score within that stage. Although gamma power was uniquely higher for rewarded trials during best performance stages, it was not a strong predictor of d-prime behavioral performance (Figure 3Fii). In contrast, alpha/beta power associated with the EXT tone continued to be strongly correlated to within session performance during the best performance stage. Together, our results demonstrated that alpha/beta oscillations emerge in mPFC in response to unrewarded trials, and that the strength of alpha/beta power during the confusion stage is a strong predictor of future task performance. In contrast, while gamma power was uniquely elevated for rewarded trials, it was not a prominent predictor of behavioral performance during the confusion or best performance stages.

### mPFC neuronal spiking encodes auditory stimuli and reward outcome, and is altered during extinction learning

Having observed significant changes in mPFC LFP oscillation dynamics throughout extinction training, we next examined multiunit activity (MUA) in mPFC. We calculated the spiking rate of mPFC MUAs during the 500ms tone presentation period (0-0.5 seconds), and the subsequent period of reward availability (0.5-2 seconds after tone onset). In the baseline stage, before extinction training, we saw a robust increase in mPFC MUA responses to both conditioned tones, throughout the tone presentation duration (Figure 4Ai). This confirms previous studies that mPFC neurons are engaged by behaviorally relevant stimuli (Maren and Quirk, 2004; Chang et al., 2010; Euston et al., 2012; Hyman et al., 2012; Moorman and Aston-Jones, 2015). mPFC MUA firing rates showed a bifurcation to the two tones during extinction training in both the tone presentation period and the subsequent reward period. In particular, although mPFC firing rates did not differ between the two tones during the 0-0.5s tone presentation period in the confusion (Figure 4Aii and 4B) or best performance stages (Figure 4Aiii and 4B), we observed a significant reduction in the spiking rate for the EXT tone during the best performance stage compared to the confusion stage, suggesting a reduction in mPFC spiking activity across the learning period for the no longer rewarded tone. During the reward window (0.5-2.0s), when animals likely integrate information about the presence or absence of reward on a given trial, PFC neurons showed an augmentation for rewarded tones, and a reduction to the EXT tone, during the confusion stage (Figure 4Aii and 4C). These effects were even more pronounced during the best performance stage where mPFC neurons showed a significant reduction in firing rate to the EXT tone (Figure 4Aii and 4C).

Because we found significant differences in overall MUA firing rates, we separated mPFC MUA activity based on the location of each electrode in cingulate, prelimbic and infralimbic regions (Figure 4Di). Across these three regions of mPFC, we found that MUA activity in the cingulate and the infralimbic cortex were more responsive to the auditory tones than the prelimbic region during all three stages of training. However, when we considered spiking activity during the reward window (0.5-2.0s) the increase in firing rates for the still rewarded tone was most strongly present in infralimbic regions (Figure 4Dii-Diii), suggesting that infralimbic regions may be more sensitive in changes of value than cingulate or prelimbic areas. We further explored this by quantifying the change in firing rate across all three stages of task performance from the three regions of mPFC. We found that none of the three regions showed firing rate changes during the tone window, but that both cingulate (Figure 4Ei) and infralimbic (Figure 4Eii) showed robust increases during the reward window while the prelimbic area did not. Interestingly, these changes in firing rates between the two tones occurred earlier in infralimbic cortex than cingulate cortex, becoming significant during the confusion stage (Figure 4Eii). While cingulate cortex also showed significant differences between tones, this did not emerge until the best performance stage (Figure 4Ei). Together, these results demonstrate subregion specific mPFC activity changes to auditory stimuli during extinction learning, consistent with an essential role of infralimbic mPFC in behavioral inhibition during extinction learning (Barker et al., 2014; Gourley and Taylor, 2016).

### mPFC neuron spiking activity in cingulate and infralimbic cortex is correlated with extinction learning

To better understand if changes in mPFC firing rate observed in cingulate (Figure 5A-B) or infralimbic (Figure 5C-D) cortex were correlated with behavioral performance, we compared the d-prime behavioral performance scores during the best performance stage to the change in firing rate for each tone in either the confusion stage or the best performance stages. Interestingly, although we did not find a significant difference in firing rate during the tone window (0-0.5s) for any of the three regions, we observed a significant correlation between changes in firing rate and d-prime performance scores in cingulate cortex for the EXT tone (Figure 5Ai: red). In addition, we found significant correlations between firing rates and d-prime performance scores during the reward window (0.5-2.0s) in both cingulate cortex (Figure 5Bii) and in infralimbic cortex (Figure 5Cii and 5Dii). This relationship appeared first in infralimbic cortex during the confusion stage (Figure 5Cii: red), before it emerged in cingulate cortex during best performance stage (Figure 5Bii: blue). In the case of cingulate cortex, the relationship to tone was unique to the reward window for the REW tone. However, in infralimbic cortex, a correlation between firing rate for both the REW tone and the EXT tone were observable during the reward window (Figure 5Bii: red and blue). Together, these findings suggest that while both cingulate and infralimbic regions predict reward presence, the infralimbic region is unique in that firing rate also encodes the absence of reward during the reward window.

**Figure 5:**
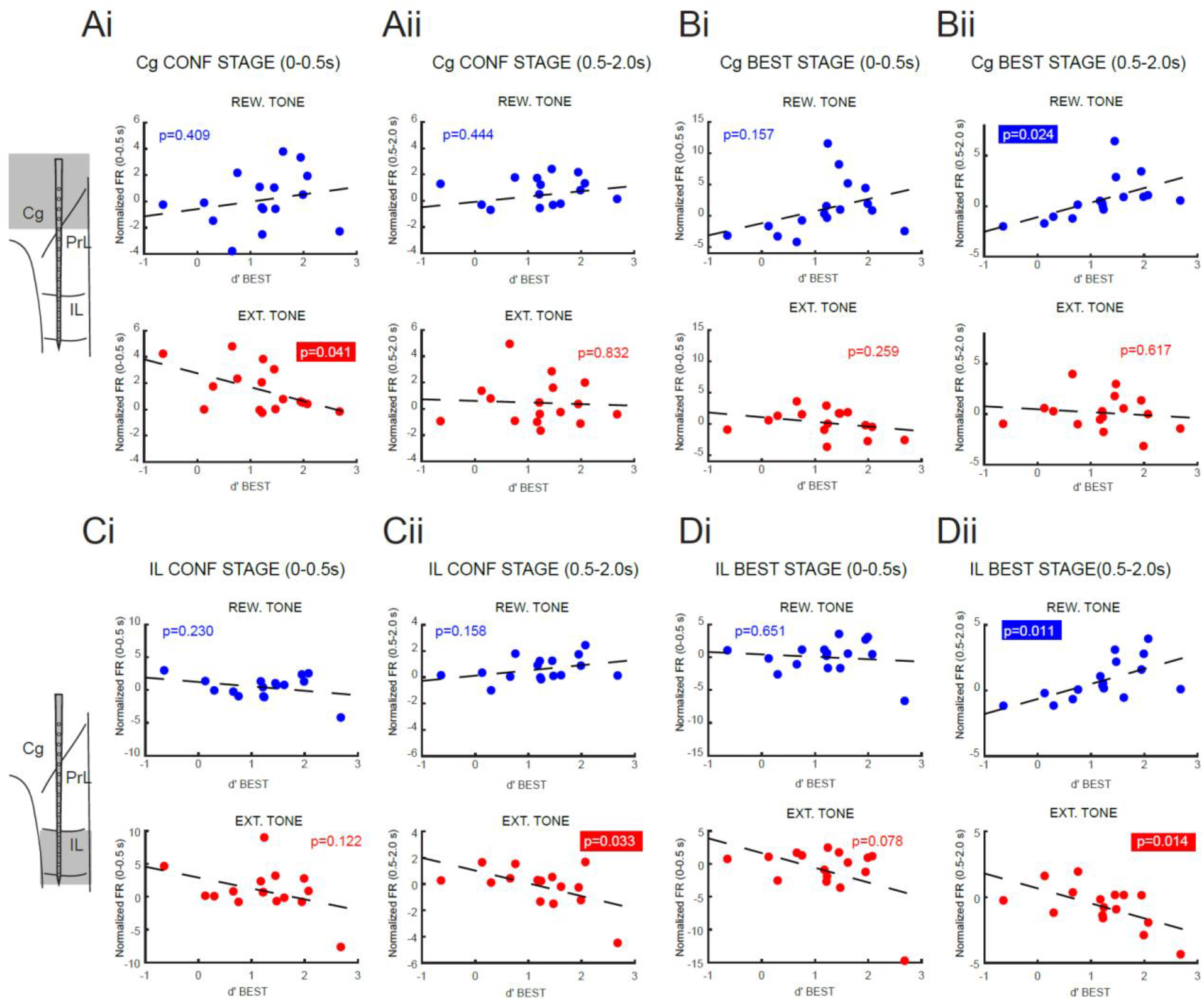
Changes in firing rate in cingulate and infralimbic cortex during the confusion stage correlates with asymptotic performance. **A:** Changes in normalized firing rate from cingulate channels during the CONF stage relative to each animal’s asymptotic BEST stage performance. Firing rate changes were compared for REW (blue:top) and EXT (red: bottom) tones for both the tone (0-0.5s) interval **(i)** and the reward (0.5-2.0s) interval **(ii)**. We found a significant correlation firing rate for the extinguished tone during the tone interval and d’ BEST performance (R=0.222, p=0.041, n = 16 mice). **B:** Changes in normalized firing rate from cingulate channels during the BEST interval relative to d’ performance. Firing rate changes were compared for both the tone (0-0.5s) interval **(i)** and the reward (0.5-2.0s) interval **(ii)**. During the BEST performance stage the strength of the FR response during the reward interval (0.5-2.0s) was strongly correlated with performance from that stage (R=0.563, p=0.023, n = 16 mice). **C:** Changes in normalized firing rate from infralimbic channels during the CONF stage relative to each animal’s asymptotic BEST stage performance. Firing rate changes were compared for REW (blue:top) and EXT (red: bottom) tones for both the tone (0-0.5s) interval **(i)** and the reward (0.5-2.0s) interval **(ii)**. We found a significant correlation between firing rate for the extinguished tone during the reward interval and d’ BEST performance (R=-0.533, p=0.033, n = 16 mice). **D:** Changes in normalized firing rate from infralimbic channels during the BEST stage relative to each animal’s performance. Firing rate changes were compared for both the tone (0-0.5s) interval **(i)** and the reward (0.5-2.0s) interval **(ii)**. During the BEST performance stage the strength of the FR response during the reward interval (0.5-2.0s) was strongly correlated with performance for both the REW tone (R=0.612, p=0.011, n = 16 mice) and for the EXT tone (R=-0.598, p=0.014, n = 16 mice). Significant differences are outlined. **Abbreviations:** Cg; Cingulate Cortex, Prl; Prelimbic cortex, IL; Infralimbic Cortex.

### AC cortex LFP activity discriminates between trial types during the best performance stage

We next considered how auditory stimuli influence LFP responses in AC at different stages across extinction training. Laminar probes were positioned perpendicular to the cortical surface in AC, and each probe consisted of four shanks with each shank containing eight electrode sites spaced 100um apart. This recording configuration allowed us to sample the majority of cortical depth in the AC and perform current source density (CSD) analysis using LFPs recorded from different electrode sites on the same shank. Auditory evoked CSDs exhibit large current sinks immediately after tone onset, in the middle layers ~300-400μm from the cortical surface and represent thalamic input (Figure 6Ai, blue color). The middle thalamic input layers also show a strong source that emerges ~75 ms after tone onset (Figure 6Ai, red color). We first identified the shank with the strongest responsivity for each tone from the baseline stage by computing the sink-source amplitude difference for all four shanks to each tone (Figure 6Aii). An example CSD profile from a representative animal is shown in Figure 6Bi-Bii where the shank with the largest sink-source transition differed between shanks A and C. Across animals, we found that the most responsive shanks differed between the tones as expected based on the ~7k difference between tone frequencies. In addition, we noted the absence of a thalamocortical sink for the non-preferred tone consistent with strong lateral suppression from regions activated by the preferred tone.

**Figure 6:**
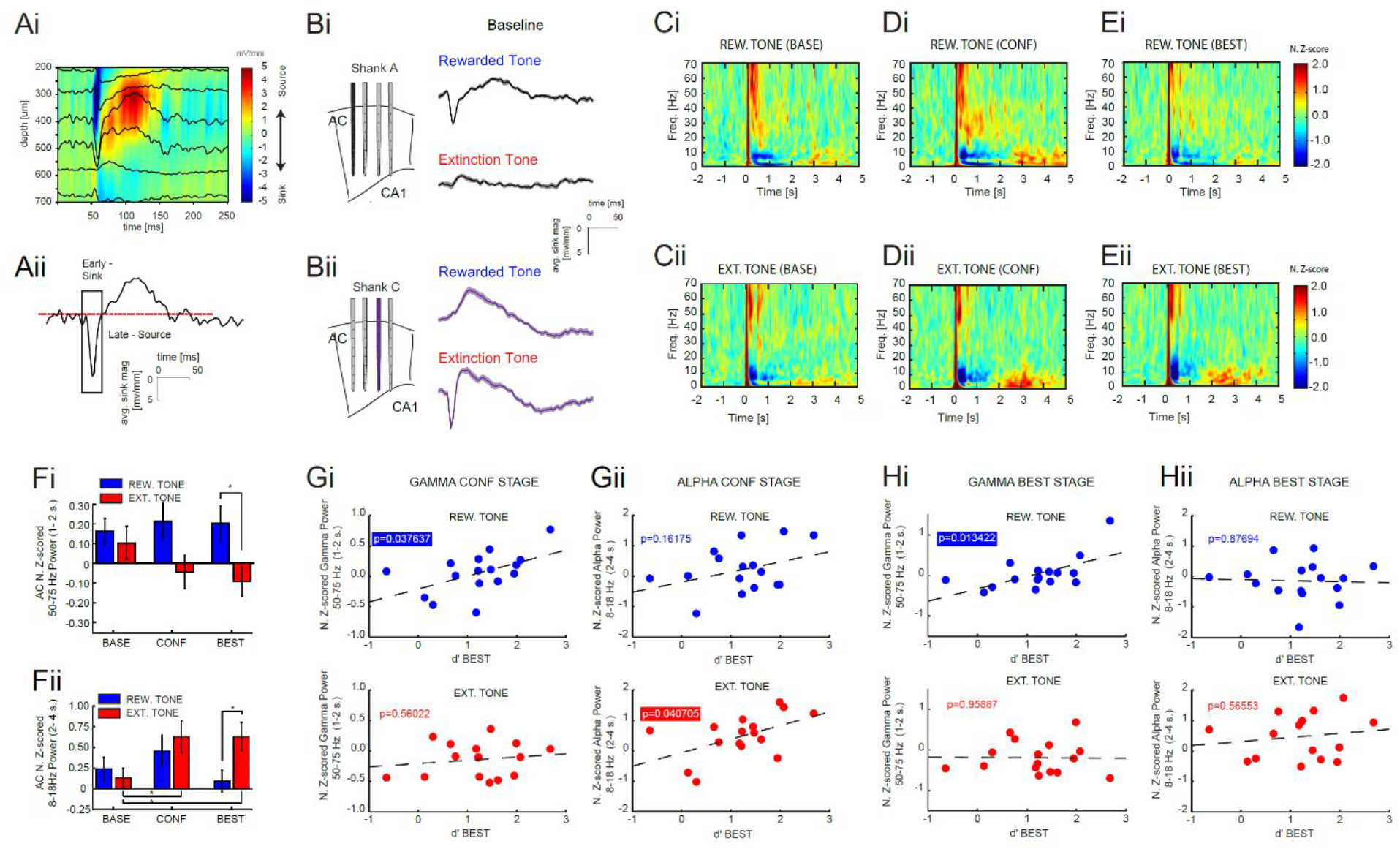
Alpha/beta and gamma power in the auditory cortex correlates with learning and is selectively altered by trial type during best performance. **A:** Tone evoked ERPs in auditory cortex overlaid on CSD calculated from laminar channels **(i)** and the sound-evoked CSD response from the channel exhibiting the largest thalamocortical response **(ii)** from a representative animal. Red dotted line represents 0 mV/mm and zero crossings represent changes in current flow. **B:** CSD response from the channel with the largest thalamocortical response for the shanks most responsive to the REW **(i)** tone (Shank A: black) and the EXT **(ii)** tone (Shank C: purple) during the baseline stage. The response to the non-preferred tone is shown for comparison and is represented by a strong source in the absence of an initial sink. Diagram illustrates recording location for each shank with Shank A having the most rostral placement and Shank D being the most caudal. **C:** Average normalized wavelet spectrogram of AC LFP for shank with the largest thalamocortical response from CSD analysis aligned to tone onset during the BASE stage for the rewarded tone **(i)** and the extinguished tone **(ii). D,E:** Same as **C** except for the CONF **(D)** and the BEST **(E)** performance stages from all mice for the rewarded tone **(i)** and the extinguished tone **(ii)**. Tone onset begins at zero. Power was Z-score normalized to the 2s window before tone onset. Note the increase in alpha/beta power for both tones during the CONF window that persists into the BEST stage for the EXT tone during the post-reward window (2-4s). **F:** Quantification of changes in gamma power (50-75 Hz) in the 1-2 second window following tone onset **(i)** and in alpha/beta power (8-18 Hz) in the 2-4 second post-reward window across the three stages of training **(ii)**. Gamma showed a significant effect of tone (ANOVA(1,15), F=6.005, p=0.024). Post hoc analysis revealed significant differences between tones in the BEST performance stage (p=0.0109). Alpha/beta also showed a significant tone by block interaction (ANOVA(1,15), F=3.903, p=0.0312). Post hoc analysis also revealed significant differences between tones in the BEST performance stage (p=0.0391). **G:** Relationship of gamma power **(i)** and alpha/beta power **(ii)** from the CONF period against future d’ performance for each tone (blue:REW tone and red:EXT tone). There was a significant correlation between the strength of gamma power following REW tone presentation on CONF trials to asymptotic d’ performance (R=0.523, p=0.038, n = 16 mice). Additionally, although alpha/beta power was upregulated following both tones in AC, there was only a significant correlation with alpha/beta power and asymptotic d’ performance following the EXT tone (R=0.516, p=0.041, n = 16 mice). **H:** Relationship of gamma power **(i)** and alpha/beta power **(ii)** from the BEST period against d’ performance for each tone (blue:REW tone and red:EXT tone). There was a significant correlation between the strength of gamma power following REW tone presentation to d’ performance (R=0.603, p=0.013, n = 16 mice). Error bars represent S.E.M. (*=p<0.05, **=p<0.01).

After identifying the most responsive shank for each tone from the baseline (REW shank or EXT shank, respectively), we compared the changes in LFP spectrogram on that shank following tone onset, during the baseline, confusion, and best performance stages (Figure 6 C-E). Consistent with the changes noted in mPFC, tone presentation produced robust changes in the AC LFPs during the baseline stage, with no quantifiable difference between the two tones (Figure 6Ci-Cii). Sound evoked changes in LFP power at gamma frequencies increased immediately after tone onset, whereas the lower frequency (0.5-20Hz) power decreased during the reward window. The reduction in lower frequency power is consistent with the presence of an active motor licking response mimicking findings from the mPFC.

When we considered changes relative to the stages of extinction training, we noted that during the confusion stage, we saw the emergence of strong gamma frequency oscillations during the 1.0-2.0s reward period, followed by the emergence of the 8-18Hz alpha/beta oscillation in the post-reward period (2.0-4.0s), for both the REW and EXT tones (Figure 6Di-Dii). We observed that like the mPFC, this increase in alpha/beta frequency oscillations remained only for the EXT tone in the best performance stage (Figure 6Ei-Eii), suggesting that alpha/beta oscillations differentiate between task relevant stimuli in both mPFC and AC. Interestingly, changes in AC LFP alpha/beta power were not significantly different until the best performance stage (Figure 6Fi-Fii), in sharp contrast to the mPFC where alpha/beta power differences between tones were observable during the confusion stage (Figure 3C and 3D). Similar to mPFC, we found that the strength of alpha/beta during the confusion stage was also predictive of future d-prime performance during the best performance stage (Figure 3Gii). However, in contrast to mPFC, AC LFP gamma power was a strong predictor of d-prime performance during both the confusion and best performance stages (Figure 6Gi and 6Hi). In summary, these results demonstrate that both gamma and alpha/beta frequency oscillations distinguish trial types in AC during extinction training.

### Auditory cortex neural firing rates are modulated during extinction learning

To better understand whether the changes in LFP oscillations in AC were associated with changes in AC neural firing rates, we measured multiunit activity across the depth of each laminar shank. We found that both tones evoked a large auditory response at tone onset, ~8X increase compared to the baseline firing rate before tone presentation (Figure 7Ai and 7Bi). The largest increase occurred on electrode sites 300-400μm from the cortical surface, corresponding to the thalamic input layers. In addition, we found that the deeper layers of AC exhibited prolonged responses to the tones throughout the tone window, and for several hundred milliseconds after the tone ended. When we compared changes in firing rate during the confusion and best performance stages, we found that sites that were most response to the REW tone during the baseline stage (the REW shank, blue) show increased activity at tone offset that persisted for several hundreds of milliseconds, and was strongly prominent in the BEST performance stage (Figure 7Aiii). In contrast, sites that had shown the strongest activation to the extinction tone in the baseline stage, and prior to extinction (EXT tone, red shank), began to show reduced activity during the 0-0.5s tone window (Figure 7Biii). To better understand if these changes in firing rate changes were predictive of performance, we compared firing rate during the confusion and the best performance stages to the d-prime behavioral scores for each animal (Figure 7Ci-ii, 7Di-ii). Unlike mPFC, we found no significant correlations between AC firing rate during the confusion stage with future performance for both the tone window and the reward window (Figure 7Ci, ii). However, we found that during the best performance stage, reductions in firing rates for the EXT tone during the reward window (0.5-2.0s) was strongly related to animal performance. Taken together, these results suggest that neuronal activity in auditory cortex evolves during extinction learning, and that the change in auditory cortex neuronal spiking activity is strongly correlated with the strength of extinction performance.

**Figure 7:**
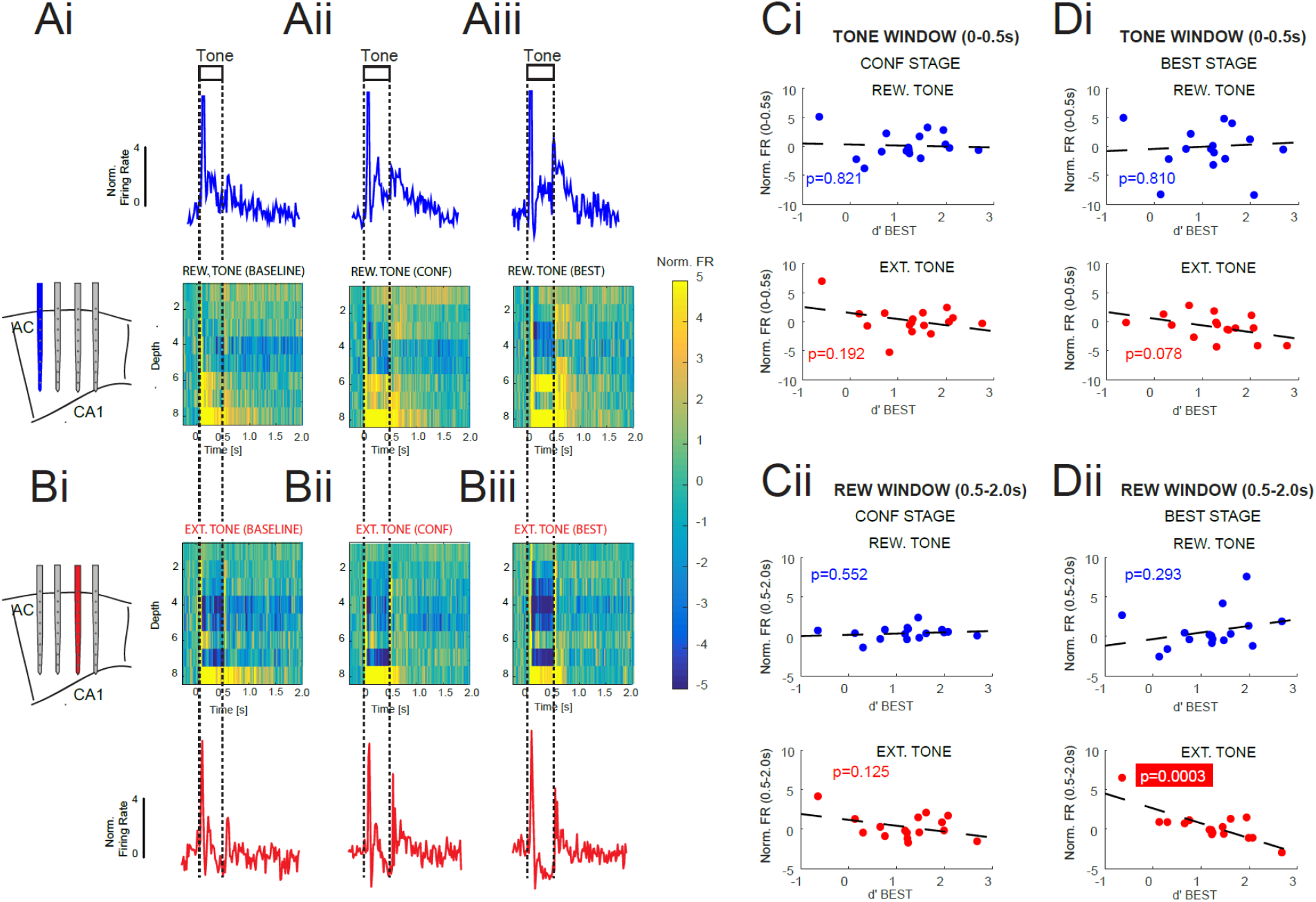
Auditory cortex neural activity is modified by extinction learning. **A:** Population firing rate plots showing activity from AC sites on the shank with the highest sensitivity to the REW tone based on current source density analysis during the BASE **(i)**, CONF **(ii)**, and BEST **(iii)** performance stages. Average sound evoked multi-unit activity (MUA) responses from all channels aligned to tone onset from all mice is shown in blue on **(top)**. The activity of all 8 channels from that shank is shown from superficial to deep across the BASE, CONF, and BEST stages in the colormap shown below. Note that during the CONF and BEST stages, neurons show more sensitivity to the REW tone following tone offset. Colormaps represent the averaged activity from all 16 mice organized by depth. **B:** Population firing rate plots showing activity from AC sites on the shank with the highest sensitivity to the EXT tone based on current source density analysis during the BASE **(i)**, CONF **(ii)**, and BEST **(iii)** performance stages. Average tone evoked multi-unit activity (MUA) responses from all channels aligned to tone onset from all mice is shown in red **(bottom)**. The activity of all 8 channels from that shank is shown from superficial to deep across the BASE, CONF, and BEST stages in the colormap shown above. Note that during the BEST period, neurons show reduced sensitivity to the EXT tone during tone presentation and no activity following tone offset in contrast to the REW tone. Colormaps represent the averaged activity from all 16 mice organized by depth. **C:** Correlation between normalized firing rate from auditory cortex channels relative to asymptotic d’ BEST stage performance in the CONF stage during the tone window **(i)**, and the reward window **(ii).** Firing rate changes were compared for REW (blue:top) and EXT (red: bottom) tones. Note we found no significant correlations between firing rate for either window during the CONF stage. **D:** Changes in normalized firing rate from auditory cortex during the BEST performance stage from the tone window **(i)**, and the reward window **(ii).** Firing rate changes were compared for REW (blue:top) and EXT (red: bottom) tones for both the tone (0-0.5s) window **(i)** and the reward (0.5-2.0s) window **(ii)**. Correlation analysis revealed a significant relationship between firing rate for the extinguished tone and performance during the reward window (R=-0.782, p=0.0003, n=16 mice). **Abbreviations:** AC; Auditory Cortex, CA1; Hippocampal area CA1.

### Coherence between mPFC infralimbic area and auditory cortex at alpha/beta frequencies during extinction learning

Given that we saw consistent changes in neural spiking activity and LFP oscillation power in both mPFC and AC during different stages of extinction training, and that these changes during confusion stage were predictive of future AC spiking and the behavioral performance during the best performance stage; we explored how these two brain regions interact by calculating LFP-LFP coherence throughout the training session. LFP-LFP coherence estimates the coordinated synchronization that is thought to reflect cortical communication (Fries et al., 2001; Buschman and Miller, 2007; Siegel et al., 2012). We measured coherence between all mPFC recording sites, and all AC sites during the baseline, confusion, and best performance stages (Figure 8A). During tone presentation (0-0.5s) and into the reward period, we found that tone presentation produced a broad increase in coherence across multiple frequency bands. We also found that raw coherence was particularly high at low theta frequencies (centered at 4Hz) before, during and after tone presentation during all three stages of training (Figure 8Ai-Aiii). To better quantify coherence changes particular to the task, we normalized coherence to the pre-tone period and considered the changes in coherence between each stage of training (Figure 8Bi-Biii). This normalization revealed enhanced coherence in the 8-18Hz frequency range in the post-reward window that lasted for several seconds after tone offset, in both the confusion and best performance stages (Figure 8Bii-Biii).

**Figure 8:**
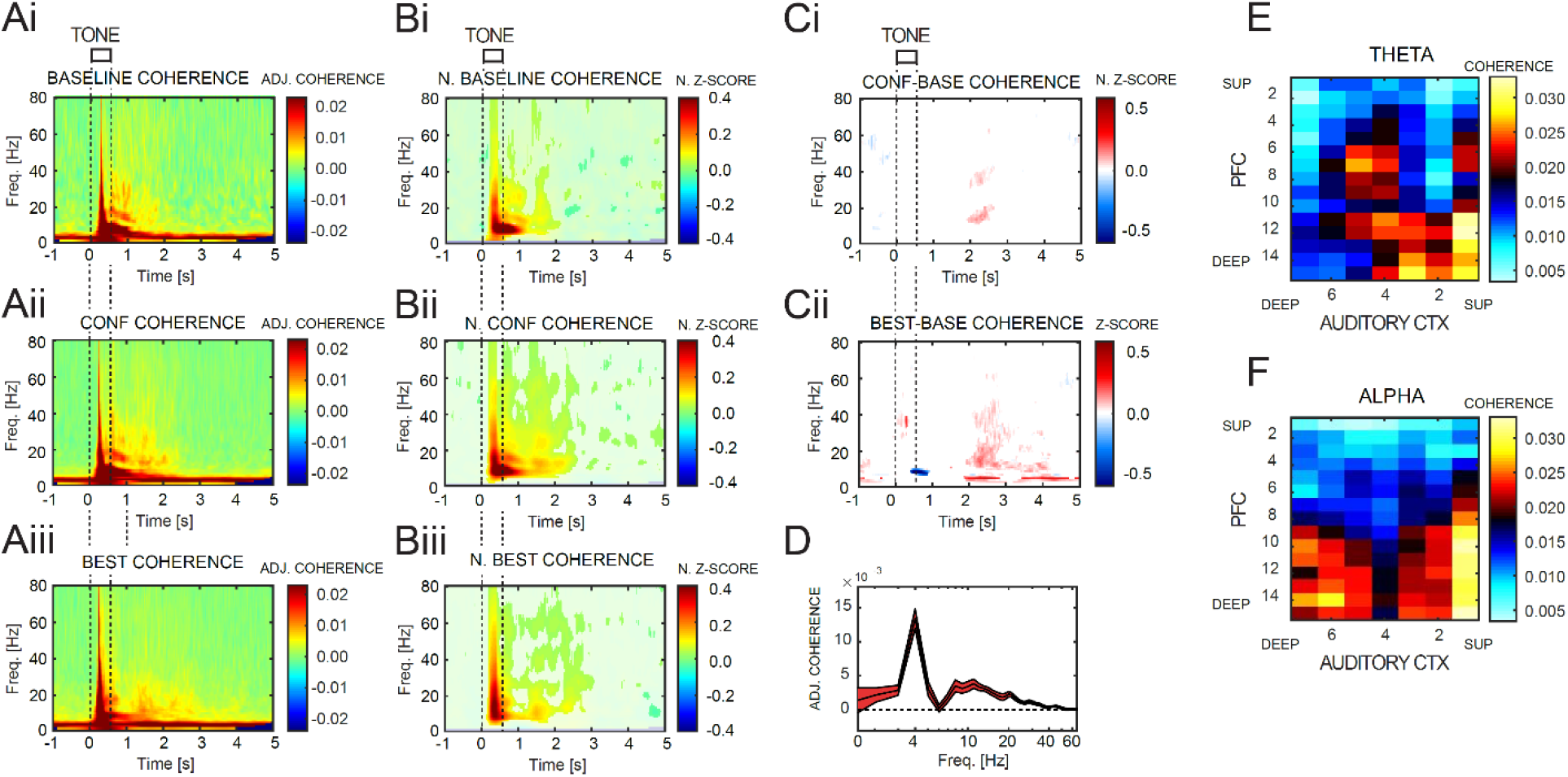
Coherence between PFC and auditory cortex is augmented at alpha/beta frequencies by extinction learning. **A:** Average adjusted coherence between the most responsive AC shank electrode sites and PFC sites aligned to tone onset during the BASE **(i)**, CONF **(ii)**, and BEST **(iii)** performance stages. Note a persistent high level of coherence at theta frequencies (3-8Hz) throughout the recording period both before and after tone and that coherence is broadly enhanced across multiple frequencies at tone onset. **B:** Normalized z-scored coherence aligned to tone onset during the BASE **(i)**, CONF **(ii)**, and BEST **(iii)** performance stages from channels shown in **(A).** Power was Z-score normalized to the 2s window before tone onset. Note the enhanced coherence following the tone at alpha frequencies (8-18Hz) during the CONF **(ii)** and BEST performance stages **(iii)** relative to the baseline stage **(i)**. **C:** Difference in coherence between the baseline and confusion stage **(i)** and the baseline and best performance stage **(ii)**. Bonferroni-corrected statistically significant differences in coherence strength are color coded (p=3e-6, n=16 mice). **(D)** Difference in adjusted coherence between the baseline and best performance stage during the post-reward window for all mice (2.0-4.0s). Note the increase in theta power and alpha power during this interval. **(E)** Matrix of absolute theta coherence (4Hz) during the post-reward window (2.0-4.0s) separated by channel during the BEST performance stage from all mice. Note the strongest theta coherence occurs between deep PFC and superficial AC channels. **F:** Matrix of absolute alpha/beta coherence (8-18Hz) during the post-reward window (2.0-4.0s) separated by channel during the BEST performance stage from all mice. Note the strongest alpha/beta coherence occurs between both superficial and deep AC regions and deep (infralimbic) regions of the PFC.

We further compared differences between stages by normalizing the coherence during the confusion stage (Figure 8Ci: difference between Bii and Bi) and the best performance stage to the baseline stage (Figure 8Cii: difference between Biii and Bi). During the confusion stage, there was a small increase in coherence unique to the post-reward window (2-4s) that emerged approximately 2 seconds after tone onset (Figure 8Ci). However, during the best performance stage, coherence between mPFC and AC was significantly elevated at both theta (4Hz) and alpha/beta (8-18Hz) frequencies in comparison to coherence during the baseline stage (Figure 8Cii and 8D).

To better understand the interactions between subregions of mPFC (cingulate, prelimbic, infralimbic), and different layers of AC (superficial, middle and deep layers), we compared coherence across all electrode sites. We found that superficial AC sites and deeper infralimbic mPFC regions showed the most elevated coherence at theta frequencies (Figure 8E). In comparison, both superficial and deep layers of AC were strongly coherent with infralimbic cortex at alpha/beta frequencies (Figure 8F), consistent with our finding of robust increase in both spiking and LFP power in infralimbic mPFC cortex for both the REW and EXT tones during the best performance stage.

## Discussion

While a great deal of progress has been made in understanding how prefrontal regions are recruited by auditory stimuli that are relevant during behavior (Maren and Quirk, 2004; Chang et al., 2010; Euston et al., 2012; Hyman et al., 2012; Moorman and Aston-Jones, 2015), far less is known about how the PFC contributes to sensory modulation as task-rules change. In particular, the process by which executive regions influence sensory regions, and the time course of their interactions. To better understand the process by which prefrontal cortex interacts with sensory cortices during stimulus processing as rules associated with goal-directed behavior are altered, we performed simultaneous recordings from AC and mPFC during an extinction learning task.

We compared LFP spectral components, multiunit firing activity, and coherence between AC and mPFC, during different stages of extinction training. Specifically, we assessed neural activity prior to extinction learning when both tones were rewarded (baseline stage), the collection of trials most immediate to where one tone is no longer rewarded (confusion stage), and the period of performance where maximal performance was achieved (best performance stage). We found that tone presentation and behavioral responses elicited broad changes in LFP oscillations that differed at each stage of the extinction task. Power at gamma frequencies was primarily associated with the reward window (1-2 seconds after tone onset) and gamma oscillations were unique to the rewarded tone during the confusion stage in mPFC (Figure 3C), whereas gamma power in AC did not differentiate the two tones until the best performance stage (Figure 6Fi). This suggests that mPFC gamma frequency oscillations may be involved in reporting the presence of reward or contribute to attentional control for relevant cues during extinction learning. Increases in PFC gamma rhythms (30-100 Hz) have been shown to be recruited as a property of long range synchrony during selective attention (Buschman and Miller, 2007; Gregoriou et al., 2009; Hipp et al., 2011). It is possibly that this process has a role in the modulation of gain during sensory processing.

Interestingly, there was also a strong alpha/beta band activity (8-18Hz) that emerged in the post-reward window (2.0-4.0s), during extinction training in both AC and mFPC. Similar to the gamma findings, alpha/beta oscillations differentiated the tones in mPFC earlier during the confusion stage before eventually differentiating the tones in AC during the best performance stage. These results suggest that both PFC and AC exhibit specific changes at alpha/beta frequencies to the EXT tone alone, and that the specificity between tones associated with this frequency is present in mPFC first during the confusion stage, before it is present in AC at the best performance stage. Alpha band (8-13Hz) activity has been linked to internal working memory processing (Jensen et al., 2002; Bollimunta et al., 2008; Jensen and Bonnefond, 2013) and thus one explanation for the increase in alpha/beta LFP power during extinction learning is that it represents working memory processes related to updating internal representation of task rules as they change. In this regard, this increase in alpha/beta power may represent an error prediction signal. Alternatively, it could facilitate the suppression of previous cue-guided behavior and the maintenance of the stimulus cue during the reward window. This interpretation is supported by recent human studies where the emergence of alpha activity acts as a suppression mechanism, to prevent initial involuntary capture by otherwise distracting inputs (van Diepen et al., 2016; de Vries et al., 2019).

We found that the emergence of gamma and alpha/beta oscillations in mPFC and AC also corresponded to periods of robust changes in spiking rates in both regions. Changes in spiking activity were observable earlier in mPFC, similar to the emergence of LFP changes. In addition, we found that the presence of alpha/beta oscillations in mPFC coincided with reduced neural spiking in infralimbic and cingulate mPFC regions, and there was a strong predictive relationship between reduced firing rate during the confusion stage and future behavioral performance during the best performance stage. We observed that mPFC neurons showed preference for reward most notably in infralimbic mPFC regions, consistent with the findings of others related to reward-seeking behaviors (Moorman et al., 2014; Moorman and Aston-Jones, 2015). In contrast to that seen in mPFC, we found that the relationship between spiking activity and behavioral performance in AC, did not emerge until the best performance stage. Our results are consistent with the findings that rapid and reversible task dependent changes in the receptive field properties of auditory cortex neurons can be sculpted by task demands (Fritz et al., 2003; Fritz et al., 2007) or through experience-dependent plasticity (Recanzone et al., 1993; Chavez et al., 2009; Winkowski et al., 2013). In particular, it has been noted that auditory evoked LFP gamma power in AC predicts associative learning and the relative level of auditory remapping during task acquisition (Headley and Weinberger, 2011). This suggests that mechanisms of attentional control may play an important role in the changes noted during extinction learning.

The mechanisms by which gamma and alpha/beta oscillations emerge during extinction training require further investigation. Neuromodulators including noradrenaline, dopamine, and acetylcholine are well situated to play a role in the process of updating task rules and the generation of LFP oscillation rhythms that may contribute to this process. Recent modeling experiments in humans suggest that all three neurotransmitters can contribute to learning in the face of uncertainty (Marshall et al., 2016). In particular, dopamine may contribute to rapid adaptation, whereas noradrenaline and acetylcholine contribute to unexpectedness or violations that arise from contextual changes (Yu and Dayan, 2005; Marshall et al., 2016). Further evidence suggests that cholinergic release can enhance local circuit synchrony in sensory cortices in the beta and gamma bands allowing complex, dynamic interactions to occur between top down and bottom up inputs (Kawaguchi, 1997; Xiang et al., 1998; Roopun et al., 2010; Lee et al., 2013; James et al., 2019). We further found that these changes in alpha/beta and gamma frequencies coincide with changes in auditory cortex spiking. Cholinergic involvement in auditory cortical plasticity has been well characterized previously (Quirk et al., 1997; Ji et al., 2005; Weinberger, 2007a, b) including evidence that behavioral training or basal forebrain stimulation can significantly reorganize receptive fields or temporal response profiles in auditory cortex (Kilgard and Merzenich, 1998a, b; Polley et al., 2006; Fritz et al., 2007; Froemke et al., 2007). An important future direction in understanding uncertainty learning will be to dissociate the contributions of individual neuromodulators including dopamine, acetylcholine, and norepinephrine and their interactions in contributing to this process.

Our findings are supportive of alpha/beta rhythms in mPFC contributing to neuronal and behavioral suppression. Alpha rhythms have been shown to be involved in suppressing neural responses to distracting stimuli and suppressing motor commands, as well as contributing to internal working memory processes (Palva and Palva, 2007; Mirpour and Bisley, 2013; Pani et al., 2014). The process by which these rhythms might influence action is not currently known, although direct cortical striatal projection manipulation can bias activity during sensory discrimination tasks (Znamenskiy and Zador, 2013). The change in alpha/beta power coincided with a reduction in neural spiking responsivity for the EXT tone, and enhanced coherence between superficial auditory cortex and infralimbic regions of the mPFC during the best performance stage. This is consistent with observations that long-range, coordinated synchronization between top-down processing regions and bottom-up sensory regions is an inherent component of performance in discrimination tasks, and that they are important for the maintenance of newly learned associations, and/or contribute to updating expected value when outcome contingencies are altered.

## Materials and Methods

All procedures involving animals were approved by the Boston University Institutional Animal Care and Use Committee (IACUC). A total of 16 mice (3-6 months-old on the day of recording) were used in this study. Original breeding pairs of Chat-Cre(B6;129S6-Chat^tm1(cre)Lowl^/J) were obtained from Jackson Laboratory, (Maine) and all breeding was done in house.

### Surgical Procedures

Mice were surgically implanted with a head-plate as described previously (James et al., 2019). Briefly, under isoflurane anesthesia, a custom head-plate designed to allow access to PFC and AC was anchored to the skull with 3 stainless steel screws. A fourth screw was connected to a metal pin and placed in the skull above contralateral parietal cortex to serve as the ground.

### Acclimation and Behavioral Training

Mice were gradually titrated down to a single hour of water access over 7 days while undergoing gentle handling before beginning training on a simple operant association task. Following water deprivation, animals were conditioned to being restrained while having access to a retrieval port for water reward. Water reward consisted of a 5μL droplet of 0.02% saccharine water and was delivered on an FI8 second schedule for 2-3 days. Once animals collected in excess of 200 rewards over a 60 min training session they moved to the Pavlovian training phase. In the Pavlovian phase animals were introduced to the two (0.5s) conditioned stimulus (CS) tones (3075 Hz and 10035 Hz) presented at 75dB and a variable inter-trial interval (ITI: 9±3s). Tones were delivered at random coincident with free water release. Pavlovian conditioning occurred for 1-2 days before animals were transitioned to the operant version of the task. In this final version, reward delivery was now contingent on animals licking the reward port within 2s of tone onset. Mice were trained under these conditions until they responded to more than 80% of tone presentations for both tones over a 60 min session for 3 consecutive days (~350-400 trials). Training under this condition typically took 5-7 days. After this criterion was met, mice underwent a sham recording session in order to habituate them to the events and extra procedures performed on the recording day. Briefly, mice while under extra lighting had the majority of their bone cement removed from the skull and were immobilized for an additional hour before starting training.

### Experimental Protocol: Extinction Recording Day

The first craniotomy was centered above the frontal recording site (2mm before bregma, 400 um lateral of the midline). The second craniotomy was over the auditory cortex (2.3-3.6 mm after bregma, 4-6mm lateral). With the use of two motorized micromanipulators, a multi-channel depth probe was inserted in the medial frontal lobe to a depth of 2.75mm and another multi-channel depth probe was inserted into the auditory cortex, to a depth of ~1mm. After the experiment, the probes were slowly retracted from the brain tissue. At the conclusion of the experiment, mice were euthanized with an IP injection of sodium pentobarbital, followed by transcardial perfusion.

### Data Collection

All recordings were made with a Tucker Davis Technologies RZ2 recording system in an electrically shielded sound attenuation chamber. Recordings were taken with two simultaneously implanted neuronexus silicon depth probes. A 16 contact linear probe with 100μm spacing was placed in the frontal craniotomy, while a 32 channel probe (4 shanks, 8 sites per shank) with 100μm spacing between contacts and 400μm spacing between shanks was inserted into the auditory cortex, perpendicular to the cortical surface. It should be noted that because of the curvature of the cortical surface, not all four of the shanks could be placed at precisely the same depth during each experiment. Probes were advanced until all probe contacts were within the cortical tissue. The probe was placed so that the shanks were spaced 1.2 mm along the rostro-caudal axis of the auditory cortex. In order to detect extracellular spikes, the signal was digitized at 24414 Hz and bandpass filtered between 300-5000 Hz and thresholded. All thresholds were set by eye at the beginning of each recording session. Timestamps and waveform snippets (1.3 ms) were stored for events that exceeded threshold. Local field potentials (LFPs) were low pass filtered at 1 kHz and digitized at 3051.8 Hz.

### Histology

At the end of the experiments, all mice were anesthetized and transcardially perfused for histological verification of electrode placement using cresyl acetate (C-1893; SigmaAldrich, Natick MA). Mice were perfused with 30 mL buffered saline, followed by 30 mL 4% paraformaldehyde. Brains were carefully removed and post-fixed overnight in 4% paraformaldehyde before being transferred to a 30% sucrose solution, and then sectioned coronally in 40μm slices with a freezing microtome (CM 2000R; Leica). Tissue sections were collected throughout the basal forebrain, AC, and PFC. Alternate sections were then rinsed with 0.05M PBS buffer and mounted on gelatin-coated slides and allowed to dry overnight. Sections were then rehydrated by a descending series of alcohol rinses (2 min each in 100%, 95%, 90%, 70%, and 50%) before being placed in deionized water for 5 min. Following rehydration, sections were incubated in 0.1% cresyl acetate for 5 min, followed by dehydration in an ascending series of alcohol rinses (50%, 70%, 90%, 95%, 100% (2X) for 3 min each and cleared with xylene (534056; SigmaAldrich, Natick MA) for 15 minutes. Slides were then coverslipped using DPX (06522; SigmaAldrich, Natick MA) mounting medium and analyzed.

### Data Analysis

All data analysis was performed with custom Matlab functions (MathWorks, Natick, MA). Statistical tests were repeated measure ANOVAs, for comparisons of response to tone and experiment interval, multiple comparisons between tone and experiment interval if the ANOVA showed a significant factor or interaction, and t-statistics to compute the correlation between cortical response and asymptomatic performance.

LFPs were first bandpass filtered between 1 Hz and 150 Hz, and then down-sampled by a factor of 8 (381 Hz) prior to further analysis. In awake head fixed conditions, we occasionally found motion induced artifact. To remove trials contaminated with motion artifact from subsequent analysis, we took the root mean square (RMS) in 100ms bins for each trial and rejected trials if the averaged RMS exceeded 5 standard deviations at any time throughout a trial.

### Behavioral analysis

To measure behavioral performance during the extinction task, a sensitivity index (d-prime) was used. Based on signal theory, d-prime was calculated as the difference between the z-scored hit rate of 50 successive trials for the rewarded tone and the z-scored false alarm rate, defined as the rate of responses to 50 successive presentations of the extinction tone. The first 100 trials after the introduction of the extinction task was deemed as the confusion stage. For each animal, the sensitivity index was calculated throughout the extinction task using a sliding window. The set of 100 trials following the confusion stage that yielded the largest d-prime was deemed as the period of best performance.

To further show that subjects changed their behavior during the extinction task, the average lick rate trace throughout a trial was calculated using bins 50ms long. Lick rates were then normalized by subtracting the average rate before tone onset from the overall trace. For comparisons of tone and interval, lick rates were further normalized by the lick rate during the reward window to calculate the change from baseline for each tone. For all 16 animals, 50 randomly-chosen licks were randomly selected from a portion of the inter-trial interval (Figure S1). Licks must have occurred between five and seven seconds after the onset of the last-played tone. Additionally, licks that were within three seconds of each other were excluded from selection.

### LFP spectral analysis

We used Matlab’s continuous wavelet transform with a complex Morlet wavelet as the basis function to compute spectrograms. Single trial spectrograms were first calculated, and then averaged across trials. To examine the effects of extinction learning on cortical rhythms, we normalized spectrograms by converting the average power during each time window to z-scores based on the mean power during spontaneous activity, defined as a 3 second window immediately preceding stimulus onset. Time-frequency windows of interest were quantified by averaging all normalized power within given frequency and time ranges.

To quantify the change in cortical rhythms in both the pre-frontal cortex and auditory cortex during extinction learning, two time-frequency windows of interest were used. Because tone presentation creates a broadband increase in spectral power at all frequencies, the tone period and the immediate 500ms following the tone were not analyzed. High gamma-frequency power (50-75 Hz) from 1 to 2 seconds after stimulus onset and low-frequency alpha and beta power (8-18 Hz) from 2 to 4 seconds after sound onset were analyzed. Both windows were chosen through visual inspection of the mean spectrograms in the pre-frontal cortex during confusion and best performance stages. For each time-frequency window, mean power was compared between tones and experiment stages using repeated measures ANOVA analysis. To quantify correlations between cortical rhythms and asymptomatic performance, mean power during the baseline task was subtracted from mean power during confusion and best performance for each subject. The correlation between normalized mean power and sensitivity index, hit rate, and false alarm rate were separately calculated using linear regression. T-statistics were used to determine the significance of the correlation between power and performance.

### Current source density (CSD) analysis

CSD analysis estimates the second spatial derivative of LFP signals to determine the relative current across the cortical laminar depth. CSDs were calculated using LFPs recorded from the four laminar probes inserted in AC, as described previously (Nicholson and Freeman, 1975; Mitzdorf, 1985; James et al., 2019). We averaged LFPs recorded from each electrode contact across all trials prior to CSD calculation.

First, we applied spatial smoothing across the eight electrode contacts within the same shank as described in (Sakata and Harris, 2009).

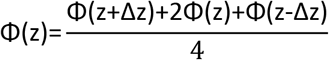

Where z is the depth perpendicular to the cortical surface, Δz is the electrode spacing, and Φ is the potential. Then we estimated the CSD as described in (Nicholson and Freeman, 1975; Mitzdorf, 1985).

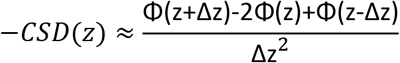

For display purposes, CSDs were interpolated linearly and plotted as pseudocolor images.

CSD analysis was also used to determine the granular layer on each shank in AC. Along each shank, the channel that had the largest sink within 50 ms of tone onset was considered to be in the granular layer. This layer identification was used to select granular layer channels for ERP latency and used for anatomical classification of laminar CSD magnitudes and MUA firing rates.

### Coherence analysis

For both auditory and prefrontal areas, their LFPs were locally referenced by subtracting each electrode contact with the mean of all contacts. We then computed the wavelet complex spectrum (Baillet et al., 2011) using morlet wavlets (6 cycles). As a coherence measure we used the Phase-locking value (Lachaux et al., 1999) defined as:

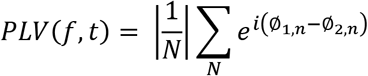

Because animals had different number of trials per condition, we adjusted the PLV value using the following equation (Aydore et al., 2013), equation 11:

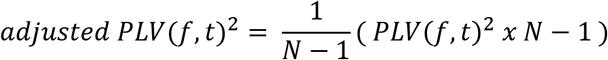

To assess statistical significance between conditions, we transformed the PLV difference into Z values (Maris and Oostenveld, 2007) and shuffled conditions labels 1000 times to obtain a null distribution. We used here the bonferroni correction as a statistical threshold.

### Spike analysis

Spike snippets were detected with manual thresholds during recording sessions and analyzed offline. Multi-unit (MU) activity was reported consistent with other auditory studies using laminar electrodes (Steinschneider et al., 2008; Guo et al., 2012; James et al., 2019). This was done because of generally lower signal amplitude obtained with laminar probes and the inability for each electrode contact to be independently positioned to best capture nearby single neuron activity leading to inconsistencies in single-unit isolation (Bragin et al., 2000). We calculated the width of spike waveforms at half of their peak-trough maximum. Spike events were only included if their width at half-maximum was between 0.1 and 0.4ms. Additionally, coincident events indicative of motion induced artifact across multiple contacts on the same probe were removed if they occurred within two samples of any other spike (83 μsec).

To analyze changes in spiking activity during extinction training, the firing rate during stimulus presentation (0 to 0.5 seconds after stimulus onset) and during the reward window after stimulus offset (0.5 to 2 seconds) were used for analysis. Spikes were binned using a 20ms time windows, and then averaged across all trials for each interval and tone combination. Then, the mean pre-stimulus firing rate was subtracted from the average firing rate trace for each interval. Similar to comparisons in LFP frequency power, repeated measures ANOVA were used to determine if the tone, interval, or interaction between the two were significant factors. Multiple comparisons were used if the repeated measures ANOVA showed a significant factor. Additionally, an N-way ANOVA analysis was done on a within-subject basis to determine significant differences in firing rate between tones within each time window.

For linear regression between spiking and performance, the firing rate during baseline was subtracted from the firing rate during confusion and best performance for each tone to determine whether a change in spiking relative to baseline correlated to asymptomatic performance.

To determine the source of any significant changes in spiking, pre-frontal cortex channels were sorted to groups based on depth. Specifically, these groups consisted of channels in the cingulate cortex (the five most superficial channels), channels in the prelimbic cortex (the middle seven channels), and channels in the infralimbic cortex (the deepest four channels). For the auditory cortex, channels were separated by their location relative to the granular layer, as identified using current source density analysis. The analysis described above was then carried out for these grouped channels.

## Acknowledgements

This study was supported by NF 1835270 and NIH 134NS111742-01 (KS and XH). X.H. also acknowledges funding from NF 1848029. We thank members of the Han Lab for suggestions on the manuscript.

## Competing Interests

The authors claim no competing financial or non-financial interests.

---

**Figure S1:**
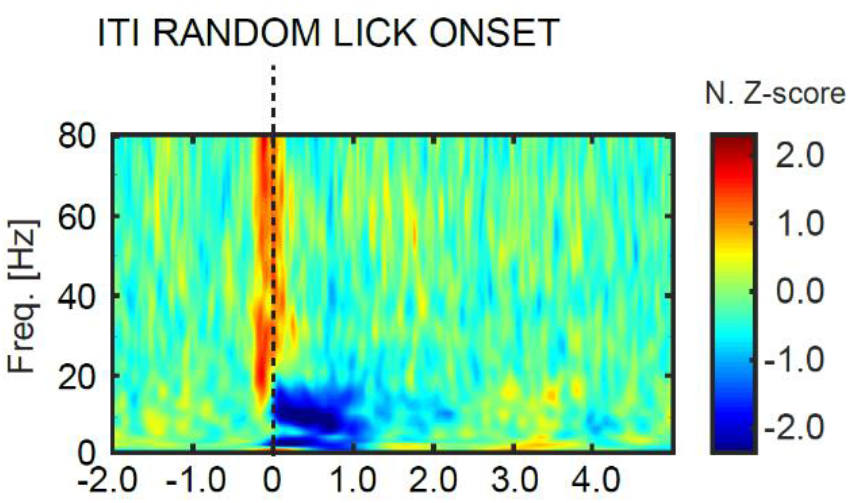
Reductions in low frequency power associated with licking behavior in the absence of reward: Average normalized wavelet spectrogram of PFC LFPs aligned to unrewarded licks randomly selected from the inter-trial interval (ITI) from all mice (n=16). Lick onset begins at zero. Power was Z-score normalized to the 2s window before lick onset. Note the reduction in low frequency power (0.5-20Hz) during the licking (motor output) period.

## Notes

### Competing Interest Statement

The authors have declared no competing interest.

